# Individual growth models support the quantification of isotope incorporation rate, trophic discrimination and their interactions

**DOI:** 10.1101/2021.05.28.446143

**Authors:** Sébastien Lefebvre, Marine Ballutaud, M. Teresa Nuche-Pascual, Sarah Nahon, Rongsong Liu, Carlos Martinez Del Rio

## Abstract

Two large but independent bodies of literature exist on two essential components of the dynamics of isotopic incorporation: the isotopic incorporation rate (λ) and the trophic discrimination factor (Δ). Understanding the magnitude of these two parameters and the factors that shape them is fundamental to interpret the results of ecological studies that rely on stable isotopes. λ scales allometrically with body mass among species and depends on growth within species. Both are often assumed to be constant and independent of each other but evidence accumulates that might be linked and to vary with growth. We built and analyzed a model (IsoDyn) that connects individual growth and isotopic incorporation of nitrogen into whole body and muscle tissues. The model can assume a variety of individual growth patterns including exponential or asymptotic growths. λ depends on the rate of body mass gains which scales allometrically with body mass. Δ is a dynamic response variable that depends partly on the ratio between fluxes of gains and losses and covaries negatively with λ. The model can be parameterized either from existing large databases of animal growth models or directly from experimental results. The model was applied to experimental results on three ectotherms and one endotherm and compared to the results of the simpler and widely used time model. IsoDyn model gave a better fit with relatively little calibration. IsoDyn clarifies and expands the interpretation of isotopic incorporation data.

## Introduction

Animal ecologists rely on stable isotope analysis (SIA) of carbon, nitrogen, and sulfur, to trace the pathways of organic matter trough food webs, to estimate trophic position, to examine intra- and inter-species trophic relationships (i.e. niche properties), to track origins and migration of animals, and to reconstruct animals’ diets (reviewed by Boecklen et al. 2011, Glibert et al. 2018 or Shipley and Matich 2020). Most of these applications hinge on the observation that the isotopic value of animal tissues resembles that of their diet with a small difference (De Niro and Eptein 1978) called trophic discrimination factor (Healy et al. 2018) and denoted by a Δ with ‰ units (see Table 1 for a list of symbols and their definitions). However, many applications of SIA in tropic ecology depend on an additional observation: the incorporation of the value of resources into consumer’s tissues after a diet change is not instantaneous, but obeys predictable temporal dynamics (Martinez del Rio and Carleton 2012). The isotopic incorporation rate (λ with units of time^-1^) is construed as the instantaneous rate of isotopic incorporation with the interpretation of 1/λ as the average retention time of an element in a tissue, and ln(2)/ λ as its half-life (Thomas et al. 2015; Vander Zanden et al. 2015).

**Table 1:** List of abbreviations and symbols used throughout the manuscript.

| Abbreviation or symbol | Unit | Definition |
| --- | --- | --- |
| $\delta$ or $\delta^H X$ | ‰ | The ratio of heavy (H) to light isotope in element X in $\delta$ notation<br>Noted $\delta$ for sake of simplicity in some occasion |
| $\Delta$ or $\Delta^H X$ | ‰ | The trophic discrimination factor i.e. the difference in $\delta$ value between the $\delta$ value of the consumer and the $\delta$ value its diet<br>Received a variety of names depending on the application including trophic shift (e.g. McCutchan et al. 2003), trophic fractionation (Vander Zanden and Rasmussen 2001), trophic enrichment factor (Post 2002) diet to tissue discrimination factor (Hussey et al. 2014). Noted $\Delta$ for sake of simplicity in some occasion |
| $\lambda$ | $d^{-1}$ | The isotopic incorporation rate also named the isotopic turnover rate with the interpretation of $1/\lambda$ as the average retention time of an element in a tissue, and $\ln(2)/\lambda$ as its half-life (Thomas et al. 2015; Vander Zanden et al. 2015) |
| $\delta_{\infty}$ | ‰ | Asymptotic value of $\delta$ in the time model of isotopic incorporation needed to calculate $\Delta$ |
| $k_g$ | $d^{-1}$ | The specific growth rate in an exponential model (also noted k).<br>Net addition of new tissues |
| $k_c$ | $d^{-1}$ | Catabolic turnover rate also noted k or m and named metabolic turnover rate. Renewal of old tissues |
| W | g | Body mass or wet weight |
| $W_{\infty}$ | g | Asymptotic body mass |
| $r_i$ | $g^{1-\beta} d^{-1}$ | Rate of body mass gains (or inputs). Named anabolic rate in von Bertalanffy growth model or Ontogenetic growth model and assimilation rate in DEB Theory. Note that underlying mechanisms may differ (Kearney 2020). |
| $r_o$ | $d^{-1}$ | Rate of body mass losses (or outputs). Named catabolic rate in von Bertalanffy growth model and maintenance rate in DEB Theory or Ontogenetic growth model. Note that underlying mechanisms may differ (Kearney 2020) |
| $\beta$ | - | Allometric coefficient |
| $\Delta_i$ | ‰ | Isotopic discrimination on the flux of body mass gains |
| $\Delta_o$ | ‰ | Isotopic discrimination on the flux of body mass losses |
| DIIM |  | Models of the dynamics of isotope incorporation in consumer tissues as a function of time or body mass. Usually one compartment first order kinetics assuming exponential growth of the consumer of which the time model (Tieszen et al. 1983; Heisslein et al. 1993) or the mass model (Carleton and Martinez Del Rio 2010) |
| DSE | - | Diet switch(ing) experiment. A controlled experiment in which a switch in diet is provoked while $\delta$ values of the consumers are measured over time and potentially body mass dynamics |
| RE | % | Relative error. Used to assess the goodness of fit of the models |

Ecologists and physiologists have conducted large numbers of experiments that describe the temporal changes of the isotopic values in consumer’s tissues after animals shift between diets of different isotopic composition (the so-called diet switching experiments, DSE, Fry and Arnold 1982, Thomas et al. 2015; Vander Zanden et al. 2015). Often, these experiments have the dual objective of estimating both λ and Δ. The results of these experiments are interpreted by fitting a family of 2 to 3 parameter models that assumes one-compartment, first order-kinetics and exponential growth of the consumers under study (e.g. Fry and Arnold 1982; Tieszen et al. 1983; Hesslein et al. 1993 and later on Carleton and Martinez Del Rio 2010). These models (referred here as isotope incorporation models, DIIM) have many virtues: they are simple, their parameters can be easily estimated and readily interpreted, and they provide an excellent fit to the temporal changes in the isotopic values of animals that follow a diet change. For instance, the widely applied time model (Tieszen et al. 1983) follows:

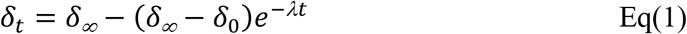

Where *δ_t_* is the isotopic composition of the consumer’s tissues over time after a diet switch, *δ_∞_* is the asymptotic value when tissues have reached steady state with the new diet (δ_d_ i.e. isotopic equilibrium) for a given incorporation rate (λ), and *δ*_0_ is the isotopic composition of the consumer’s tissues at the beginning of the DSE. Δ is often estimated as a by-product of the estimation of *δ_∞_*:

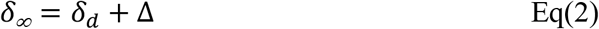

The value of λ can be partitioned into two components (Heisslein et al. 1993): the mass-specific growth rate (k_g_) that corresponds to the contribution of tissue addition due to growth and evaluated by an exponential model (with W the body mass, and W_0_ the initial body mass), and the catabolic turnover rate (k_c_) that corresponds to the replacement of existing tissue.

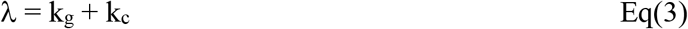

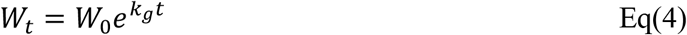

Equation 3 may not evaluate properly the contribution of k_g_ and k_c_ to λ when animal’s growth does follow an exponential pattern. For instance, MacAvoy et al. (2005) experimented on young adult mice approaching their asymptotic body mass. They observed a steady decrease in k_g_ along the course of one DSE and estimated different k_g_ values at different times. This problem can be solved by recognizing that most animals follow common asymptotic growth patterns (Kearney 2020) such as described by the von Bertalanffy growth model (von Bertalanffy 1957), the DEB theory (Kooijman, 2010) or the ontogenetic growth model (West et al. 2001) in which true exponential growth occurs only during early life stages.

Many applications of SIA in trophic ecology assume both (1) an isotopic equilibrium between the isotopic values of consumers’ tissues and its food sources (i.e. that λ is large at the time of measurement but see Marin Leal et al. 2008 and reference therein) and (2) a constant value of Δ among individuals of a population and even among different species (Phillips et al. 2014). These assumptions allows to a widely use of sophisticated user-friendly algorithms to solve mixing models that attempt to resolve a consumer’s diet composition from the isotopic values of its tissues (e.g. Parnell et al.’s (2010) Bayesian mixing model, SIAR). The importance of Δ values for the interpretation of ecological isotopic data, via mixing models, has led to compilation of large data sets of values and new methods to predict them (Healy et al. 2018). Although some tantalizing patterns between tissue type (and thus amino acid composition and isotopic incorporation), form of nitrogen excretion, nutritional status, and phylogeny have been documented (McCutchan et al. 2003; Vanderklift and Ponsard 2003; Healy et al., 2018), some of the drivers of differences in Δ remain elusive (Caut et al. 2009).

More recently, λ has received new attention (as reviewed by Carter et al. 2019). The estimation of λ is crucial to determine the time window over which diet could be reconstructed (Dalerum and Angerbjorn 2005; Phillips et al. 2014) or to model ontogenetic diet shift (Hertz et al. 2016). Incorporation rate is also a component explaining part of the isotopic variance used to evaluate the trophic niche (Fink et al. 2012; Yeakel et al. 2016). In fact, λ is a function of the body size and is expected to vary allometrically within species (Martinez Del Rio et al. 2009). This expectation has been proven to be correct between species (Thomas et al. 2015; Vander Zanden et al. 2015), even though the relationship between λ and body mass has large residual variation that remains unexplained. Assuming isotopic equilibrium and a constant Δ in isotopic ecology are possibly the result of our still incomplete understanding of the factors that shape their values: these strong assumptions should be relaxed.

Isotope ecologists have compiled large data sets of λ and Δ values estimated using DSE interpreted with first-order one-compartment models (Eq 1, 2 and 3), and assuming that these two parameters are independent and constant over the course of one DSE. Nonetheless, theoretical and empirical evidences suggest λ and Δ are dynamic and linked. On the empirical side, Lefebvre and Dubois (2016) and Gorokhova (2018) documented strong negative relationships between Δ^15^N and k_g_ (which is the dominant determinant factor of λ in rapidly growing organism, Hesslein et al. 1993) in exponentially growing animals. Villamarin et al. (2018) documented a negative relationship between Δ^15^N of crocodiles and their k_g_ that could not be accounted for a change in diet. On the theoretical side, the models of Olive et al. (2003) and Martinez del Rio and Wolf (2005) suggest a decreasing relationship between Δ^15^N and λ. The generality of this result is unknown because both Olive et al. (1999) and Martinez del Rio and Wolf (2005) modelled only the special case of animals growing exponentially. Pecquerie et al. (2010) constructed a more general approach (which is called Dynamic Isotopic budgets DIB) that assume asymptotic growth. This approach combines an accounting of the fate of different elements on the body compartments defined by the dynamic energy budget theory (DEB, Kooijman 2010). Emmery al. (2011) applied Pequerie et al. (2010)’s DIB approach to Pacific oysters (*Crassostrea gigas*) and found that Δ^15^N declined from 5 to 2‰ with increasing k_g_. Like DEB-dependent growth models, DIB models are species-specific and each case requires calibration with a high number of parameters (over 22 in the case of Emmery et al.’s (2011) application), although DEB models can be potentially parameterized with values from a huge database (Marques et al., 2018). Moreover, their application involves computationally intensive calibration and expertise in DEB that is not common among ecologists. We venture that for this reason, Pecquerie et al.’s (2010) model has not been applied widely. As far as we know, Emmery et al. (2011)’s study is its only empirical application to stable isotope studies.

Our purpose is to construct a relatively simple mathematical model that allows researching the interplay between growth, Δ and λ at the whole body and element levels while including other previous conventional models as special cases (Fig. 1). Our model permits exploring the hypothesis that λ and Δ are neither constant nor species specific, but predictably variable among individuals and dynamically linked. We hypothesize that such model would be more accurate in describing incorporation dynamics than conventional models particularly when consumer growth deviates from pure exponential trajectories (early life stages). The dual assumption of constancy and independence between λ and Δ precludes inferring their values for different life stages than the ones observed and estimated in DSEs. Another consequence of assuming static Δ and λ within the course of DSEs would be an improper estimation of their values and potentially the contribution of k_g_ and k_c_ to λ. The relative simplicity of our model facilitates its parameterization. By constructing a model that can incorporate the many ways in which animal growth has been described (for example by the ontogenetic growth model (West et al. 2001; Hou et al, 2011), von Bertalanffy growth model (von Bertalanffy 1957) and DEB theory (Kooijman 2010)), our model offers a new and dynamic perspective to interpret DSEs, but also provide a tool to help explain the still unexplained variation in Δ and λ, and in doing so provide a conceptual link between trophic isotopic ecology and the study of animal growth and life histories.

**Fig. 1.**
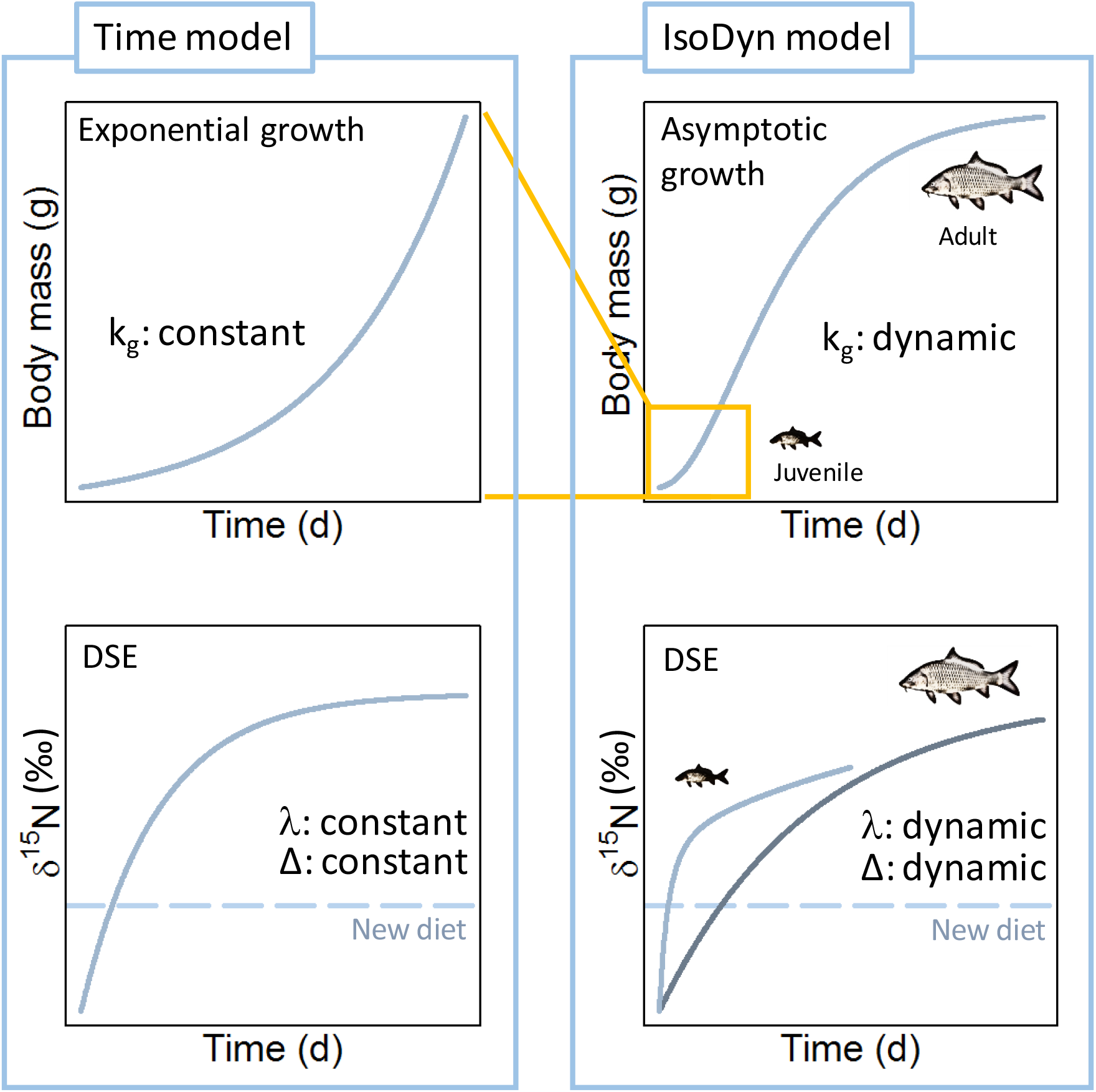
Main features of the IsoDyn model compared to the conventional time model of isotope incorporation (Hesslein et al. 1993). IsoDyn model accounts for many growth forms and offers new perspective in the interpretation of diet switch experiments (DSE) and dynamics of isotope incorporation into animal tissues in general

## Methods

### IsoDyn as a new model of isotopic incorporation

Although our model is general enough to be used by all of the stable isotopes commonly used in ecological research (C, N, S, H, and O), we will focus on nitrogen (N). The isotopic value of this element (δ^15^N) and its trophic discrimination factor (Δ^15^N) are used to estimate trophic position, and thus Δ^15^N has been relatively well studied (Post 2002; Glibert et al. 2019). Further, dietary protein is assumed to be the main driver of δ^15^N incorporation rate and Δ^15^N is less sensitive to isotopic routing than Δ^13^C (Martinez Del Rio et al., 2009).

We assume here that body mass dynamics (W_t_) of a consumer follows an asymptotic growth pattern (see supplementary material 1 for details), giving:

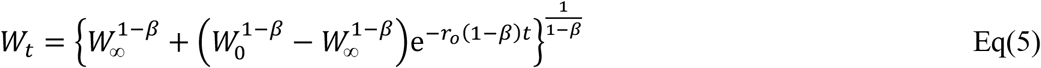

with

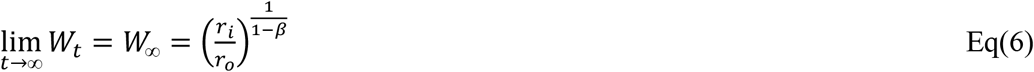

and with

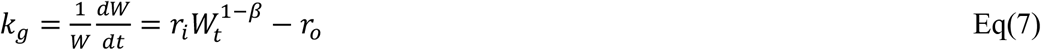

where r_i_ and r_o_ are rates of gains and losses respectively (assimilation and excretion in the case of N), β the allometric coefficient and W_∞_ is the asymptotic body mass.

Under these body mass dynamics, δ^15^N values in a consumer tissues follows (see supplementary material 1 for details):

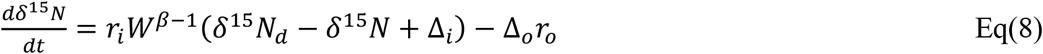

where Δ_i_ and Δ_o_ are discrimination factors on gains and losses respectively.

Eq(8) is a linear differential equation without a general analytical solution when β is lower than 1 (i.e. for asymptotic growth models such as the von Bertalanffy model). However, a discrete approximation can be done for small dt:

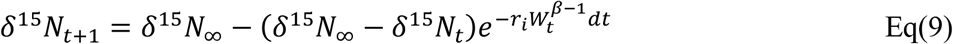

This approximation of Eq(8) then parallels the time model in Eq(1), most commonly used to describe and interpret isotopic incorporation data in DSEs, but with *δ^15^N_∞_* and λ depending on the body mass at time t:

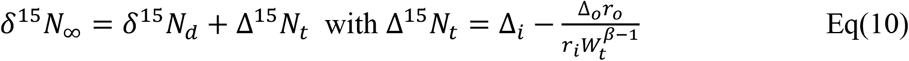

and

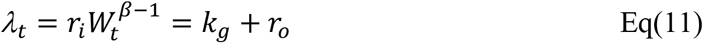

Eq(10) and eq(1) differ between them in that the terms equivalent to *δ^15^N_∞_* and λ (eq(2) and eq(3) vs eq(10) and eq(11) respectively) vary with time. Another difference is that partitioning between growth and catabolism is variable in eq(11) because k_g_ is variable while it was constant in eq(3) due to the assumption of exponential growth (eq 4). Note that r_o_ is equivalent to k_c_ or m in eq(3). These equations highlight the predictable dependence of λ and Δ^15^N on the parameters that shape growth (β, r_i_ and r_o_). A large number of studies report values for these parameters (see e.g. West et al. 2001 for the ontogenetic growth model; and Kooijman 2010 for von Bertalanffy 1957). Eq(8) also means that the isotopic incorporation rate for N can be estimated from the dynamics of body mass as long as the proportion of N in the body mass (p_N_) is constant (see supplementary material 1 for explanations). This important feature applies to any pool of element as for instance carbon or sulfur in the absence of isotopic routing.

### Simulations and parameter estimations

One of the major advantages of our model is that it can be parameterized and fitted readily with available information data or data that can be gathered in DSE. Describing dynamics of body mass require three parameters that are r_i_, r_o_ and β, and only two parameters if we consider β following a common framework (e.g. β=2/3 in the case of the von Bertalanffy growth model). Two additional parameters are needed, Δ_i_ and Δ_o_, for simulating the incorporation dynamics of stable isotopes. Simulations presented here, were generated following a numerical integration algorithm under R software v.4.0.3 (2019) using the package DeSolve (Soetaert et al. 2010) to solve eq (8). Because fitting the four parameters (r_i_, r_o_, Δ_i_ and Δ_o_) from data on isotope incorporation alone (δ^15^N values over time) is not possible, the parameters can be estimated in two ways which can be called simultaneous and sequential and by using dynamics of body mass in parallel.

In simultaneous estimation, the parametrization of the four main parameters (r_i_, r_o_, Δ_i_ and Δ_o_) can be done using both the dynamics of body mass and stable isotopes incorporation. For simplicity, we chose to perform the calibration considering that Δ_i_ and Δ_o_ as opposite values but equal in absolute value (Δ_i_ = - Δ_o_). This means that the same isotopic discrimination was applied to gains and losses, an assumption also done by Flynn et al (2018) in a mechanistic simulation model. Parameter estimations were performed using the Nelder Mead function of the lme4 package which allows to set boundaries for the parameters (r_i_ and r_o_ must be positive). The function minimizes the sum of two symmetric bounded loss functions (hereafter named the cost function) which accounts for the difference between predictions and observations for dynamics of body mass and stable isotopes respectively. This cost function is ideally suited to fit several models to several dataset (Marques et al. 2019). Local minima can be found during the optimization process. In order to ensure to detect the global minimum, the initial starting values of the parameters were randomly selected and the procedure is performed twice first (N=2). Then, the process continues (up to N=20) until the value of the cost function for the last set of parameters is lower than the best set by 5%. Parameter sets in which some parameters stuck to the boundaries were systematically deleted. Interval estimates of parameters were evaluated using a bootstrap method (N=500) by adding log-normally distributed scatter (mean coefficient of variations of observations) to the predictions with replacement of the original data sets (Marques et al. 2019). We then compare the parameters of the IsoDyn model with the time model partitioning λ into k_g_ and k_c_ (eq 1, 2, 3 and 4). The time and exponential models were fitted with the nls2 package. Goodness of fit of all the models was assessed by the relative error (RE) as calculated by Marques et al. (2019):

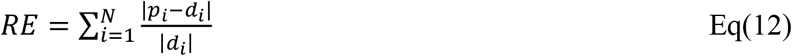

where p_i_ and d_i_ are prediction and observation, respectively, for a given data point i and N is the total number of data points.

The sequential estimation consists first in obtaining estimates for r_i_ and r_o_, which allows estimating body mass over time as well as λ. This can be done either by conducting experiments and fitting the parameters of von Bertalanffy (1957) or West et al. (2001)’s equations. In the absence of sufficient experimental data, r_i_ and r_o_ can be obtained from data bases developed from DEB theory such as “Add my Pet” (Marques et al., 2018) or the ontogenetic growth model (West et al. 2001; Hou et al. 2011) or studies on the selected species. Then, the estimation of Δ_i_ and Δ_o_ were done in a second step by implementing the values of the three previous parameters in Eq (8).

R code for all analyses, figures and tables is available from GitHub (https://github.com/Sebastien-Lefebvre/IsoDyn)

### Data sets

For our model to be calibrated or validated, dynamics of body mass in parallel to dynamics of nitrogen stable isotope incorporation are needed and these combinations are not often reported in experimental studies. Our predictions apply to the whole organism. However, it is generally assumed that muscle tissue and other structural tissues form the majority of an organism’s body mass and that consequently isotopic dynamics of the whole organism can be approximated by the ones of muscle tissues (Thomas and Crowther 2015). We have then selected four studies to highlight the different ways to estimate parameters in the context of DSEs. The first study applied on young adult mouse (*Mus musculus*) approaching the maximum body mass (MacAvoy et al., 2005). Stable isotope incorporation dynamics were measured over 112 days DSE on skeletal muscle (δ^15^N_m_) using an experimental diet. *Mus musculus* is a small endotherm species with a maximum body mass of ca 25 g. In the second study, Pacific yellowtail (*Seriola lalandi*) juveniles were used for a 98 days DSE in which incorporation dynamics of stable isotope ratios of nitrogen of dorsal muscle (δ^15^N_m_) were measured (Nuche-Pascual et al. 2018). Fish were fed with a commercial diet. *Seriola lalandi* is a large ectotherm species with a maximum body mass of ca 193 kg. In the third study, sand goby (*Pomatoschistus minutus*) late juveniles were used for a 90 days DSE and δ^15^N_m_ values were measured (Guelinckx et al. 2007). Fish were fed with a commercial diet. *Pomatoschistus minutus* is a small ectotherm species with a maximum body mass of ca 7 g. These three first studies were calibrated following the simultaneous estimation. In the fourth and last study, the growth rate of common carp (*Cyprinus carpio*) was manipulated by changing food availability providing four different diet switching experiments lasting eight weeks (Gaye-Siesseger et al. 2004). Only start and end values of body mass and δ^15^N values of the whole body (δ^15^N_b_) were originally provided for this study. *Cyprinus carpio* is a medium ectotherm species with a maximum body mass of ca 40 kg. This last study was calibrated using the sequential method for estimates of r_i_ and r_o_ of this species as described by the DEB theory (ESM 2).

## Results

Although our model shares a variety of characteristics with previous models, it has specific ones. In this section we highlight two of those: 1) IsoDyn model makes explicit links between growth and isotopic incorporations patterns; 2) the model allows parameterizing and fitting existing patterns on the dynamics of isotopic incorporation particularly when consumer growth deviates from pure exponential trajectories. Before considering our analyses on the four case studies, we considered a few general traits of our model that distinguishes it from previous ones.

### Growth and isotopic incorporation patterns are predictably linked

Our model predicts that the relationships between body mass and the isotopic value of tissues as a function of time are shaped by a set of common parameters (i.e. r_i_, r_o_ and β). Examples of such relationships and their effects on body mass, λ and δ^15^N dynamics are given in Fig. 2 for three virtual species characterized by different values and ratios of r_i_ and r_o_, a common β=2/3 and initial body mass W_0_=0.1 g. Species 1 (Sp1) and species 2 (Sp2) have the same asymptotic body mass (W_max_=64 g eq(7)) but differ in their mass gains and losses rates by half (r_i1_ = 0.2 g ^1/3^ d^-1^, r_o1_ = 0.05 d^-1^, r_i2_ = 0.1 g ^1/3^ d^-1^, r_o2_ = 0.025 d^-1^). As rate of losses (r_o_) governs the steepness at which W_max_ is reached, Sp1 reached its W_max_ faster (Fig. 2A). Species 3 has higher r_i3_=0.6 g ^1/3^ d^-1^ and r_o3_=0.2 d^-1^ but lower W_max_ = 27g. Dynamics of isotopic incorporation are then explained by two components, λ and δ^15^N_∞_ value which are both dynamic in our model (see equation 10 and 11).

**Fig. 2.**
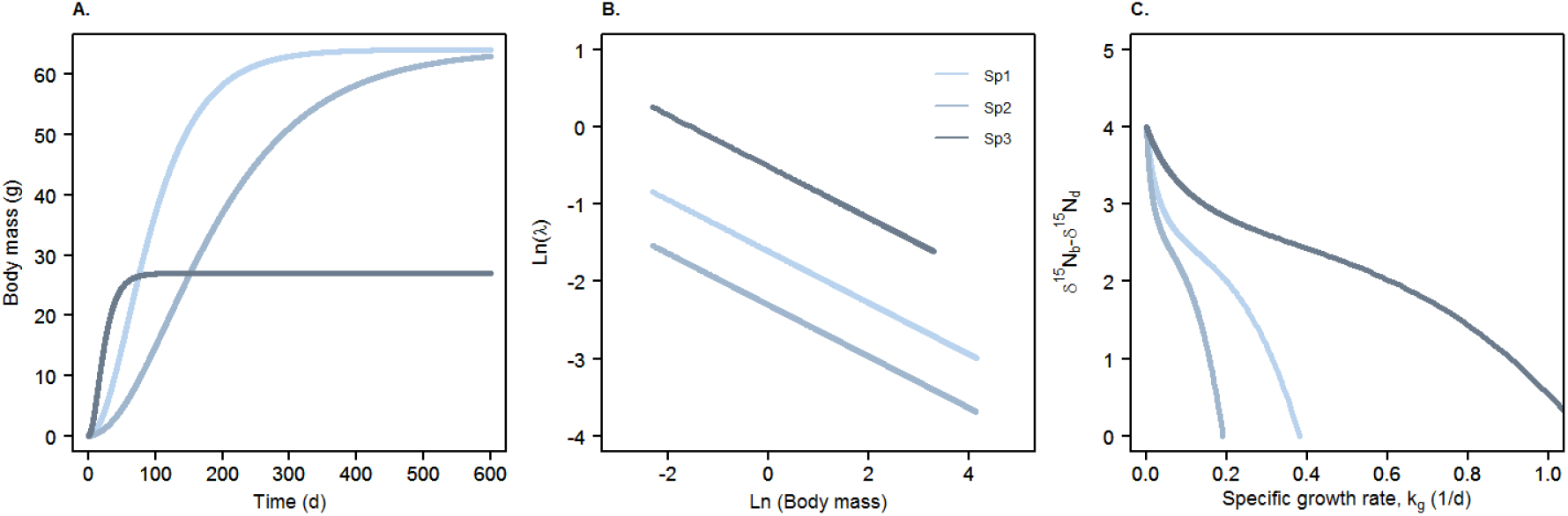
General patterns of the IsoDyn model over time for three virtual species (see text for details) with an allometric coefficient β = 2/3. The same parameters that shape growth also shape isotopic incorporation. A) Body mass over time B) Range of values of isotopic incorporation rate ln(λ) = ln(r_i_) + (β-1) ln(W), C) δ^15^N difference between body and diet as a function of specific growth rate (k_g_) with Δ_i_ = 2‰, Δ_o_ =-2‰ and δ^15^N_d_=0‰. k_g_ was calculated following eq(15)

Typically, λ decreases over time along with body mass. On a ln/ln scale, the slope is negative and equals β-1, and the intercept is ln(r_i_) (Fig. 2B); r_i_ and thus λ are higher in Sp3 than Sp1 and Sp2. The range of λ displayed by one species depends on the difference between W_0_ and W_max_, which is lower for Sp3. As λ and k_g_ decrease, δ^15^N difference between body and diet (δ^15^N_b_-δ^15^N_d_) increases (Fig. 2C). When growth approaches zero, δ^15^N dynamics are dominated by flux of body mass losses (i.e. excretion) and δ^15^N_b_-δ^15^N_d_ reaches its maximum (i.e. Δ_i_ - Δ_o_ = 4‰). On the opposite, when growth tends to its maximal value, δ^15^N_b_-δ^15^N_d_ approaches δ^15^N_d_ (here set at 0) and δ^15^N dynamics are dominated by the flux of mass gains (i.e. assimilation). The inflexion characterizes the trade-off between fluxes of gains and losses dominance in δ^15^N dynamics. Then, the range of these values depends on the extent of k_g_ performed by the species between initial body mass (i.e. birth W_0_) and W_max_.

These results suggest that the δ^15^N dynamics obtained in DSE will depend on the stage of growth at which experiments are done. Typical DSEs were performed using features of the three species above at two life stages (juveniles i.e. from W_0_ and adult at W_max_). As our analysis of special cases indicates, δ^15^N_b_-δ^15^N_d_ values will be lower in experiments involving organisms growing in the early, quasi-exponential, phases of body growth than in animals of the same species that have reached or are close to the asymptotic body mass.

In Sp1 and Sp3, adults reach the asymptotic isotopic composition faster, because λ (which equals r_o_ in adults) remains quite high even for adult while δ^15^N_b_-δ^15^N_d_ is maximum (Fig. 3). For the young of these species, although λ is even higher than for adults, δ^15^N_b_-δ^15^N_d_ is still increasing and this prevents from reaching fully the asymptote for juveniles of Sp1. The pattern is a bit different for Sp2 (Fig. 3). As juveniles of Sp2 are still performing high k_g_, then λ remains high and the δ^15^N_b_-δ^15^N_d_ although increasing, is still low: asymptotic isotopic composition remains a moving target and cannot be fully reached. In Sp2 adults, λ is twice lower than for Sp1, and its value does not reach the asymptotic value within the 100 days of the experiment.

**Fig. 3.**
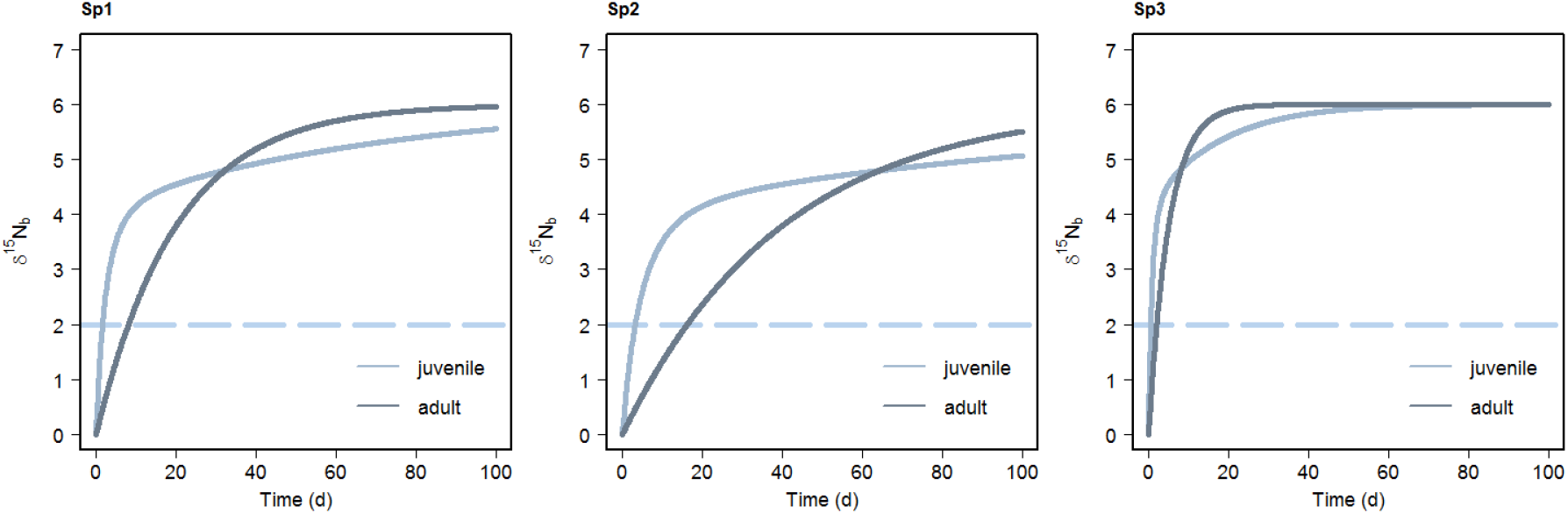
Typical 100-day Diet Switch Experiment for three species with different growth patterns (described in text and Fig. 2) and comparing patterns for juveniles and adults. Figures describe changes in δ^15^N_b_ of whole body over time for Sp1, Sp2, and Sp3. In each species, the experiment for juveniles starts at W_0_. For adults, experiments start at W_max_. The dash line represents δ^15^N_d_ value of the new diet

### Case study: calibration using the simultaneous parameter estimation

The three species studied are an endotherm (the house mouse *Mus musculus*) and two ectotherms (the two fish species Pacific yellowtail, *Seriola lalandi* and sand goby, *Pomatoschistus minutus*), at different life stages and as a consequence, in different growth situations at the time of DSE. The young adult mice approached their asymptotic body mass so that their growth rate gradually decreased during the experiment. The mouse body mass gain was about 25% (Fig. 4). Individuals of both fish species were juveniles and showed a linear body mass increase for Pacific yellowtail and an exponential increase for sand goby with a body mass gain of 600% and 100% respectively (Fig. 4).

**Fig. 4.**
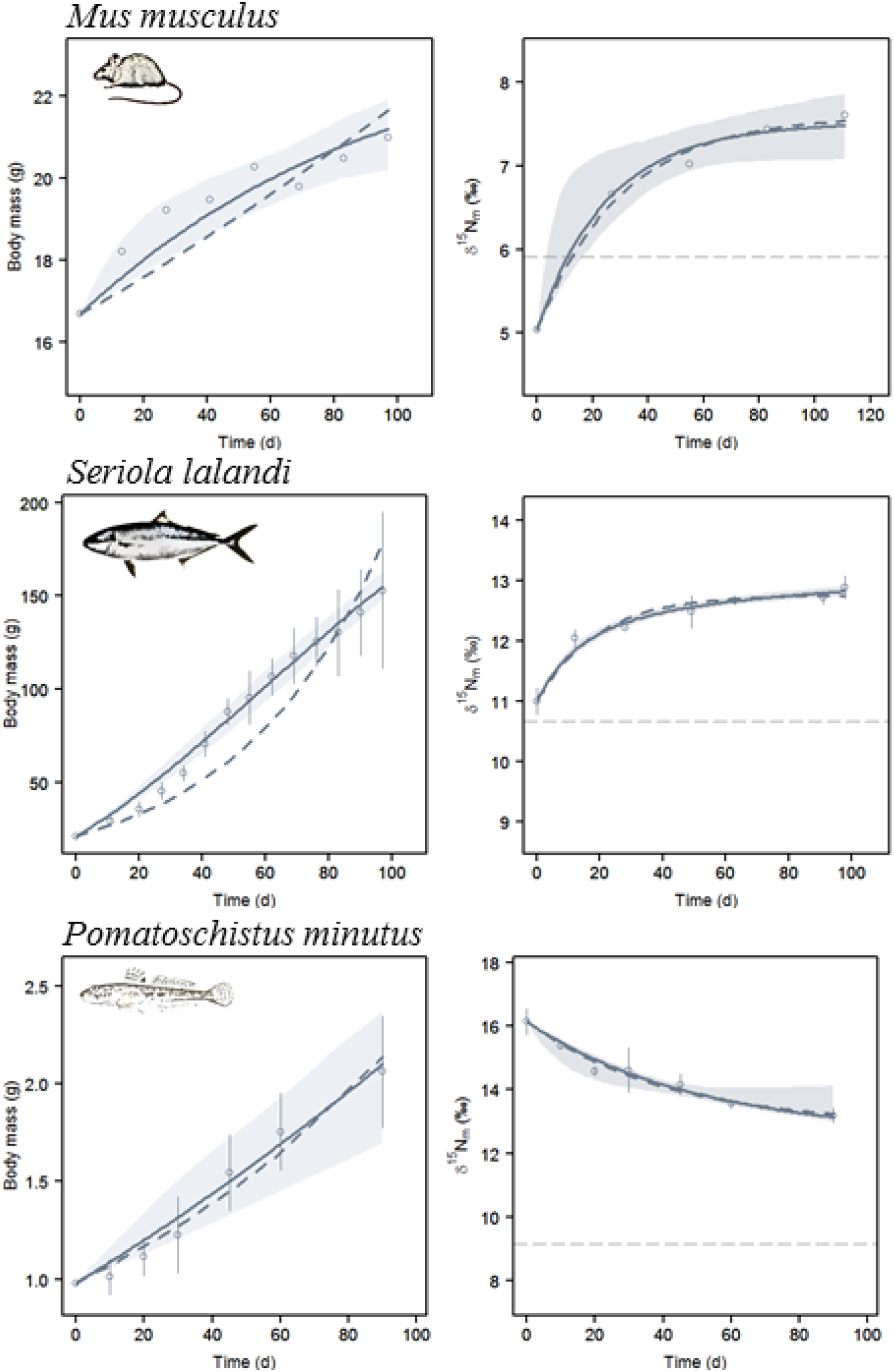
Changes in body mass (in g; left column) and δ^15^N_m_ values of muscle tissue (in ‰; right column) in three species (young adult mouse *Mus musculus* data from MacAvoy et al. 2005; Pacific yellowtail juvenile fish *Seriola lalandi* data from Nuche-Pascual et al. 2018; sand goby juvenile fish *Pomatoschistus minutus* data from Guelinckx et al. 2007). Open circles are observations (mean ± sd), solid lines are predictions from Isodyn model (eq 6 and 13), dotted lines are predictions from the time model (eq 1 and 4). Colored envelopes are 2.5 and 97.5 quantiles of IsoDyn model predictions. Grey dashed lines are δ^15^N_d_ of the new diet

A comparison between the conventional isotopic incorporation time model and IsoDyn model was done. Both models displayed very good and comparable goodness of fit concerning δ^15^N values (as estimated by the RE Table 2). However, the two models differed strongly in the prediction of body mass dynamics for two species (mouse and Pacific yellowtail) which did not follow exponential growth patterns. In these two cases, the exponential model fitted poorly to the data whereas IsoDyn model fitted better. Both models displayed a very good fit for sand goby growing exponentially. Therefore, IsoDyn model performed better than the time model regarding simultaneously the body mass and δ^15^N dynamics (mean RE Table 2) when body mass dynamics did not follow an exponential pattern.

**Table 2.**
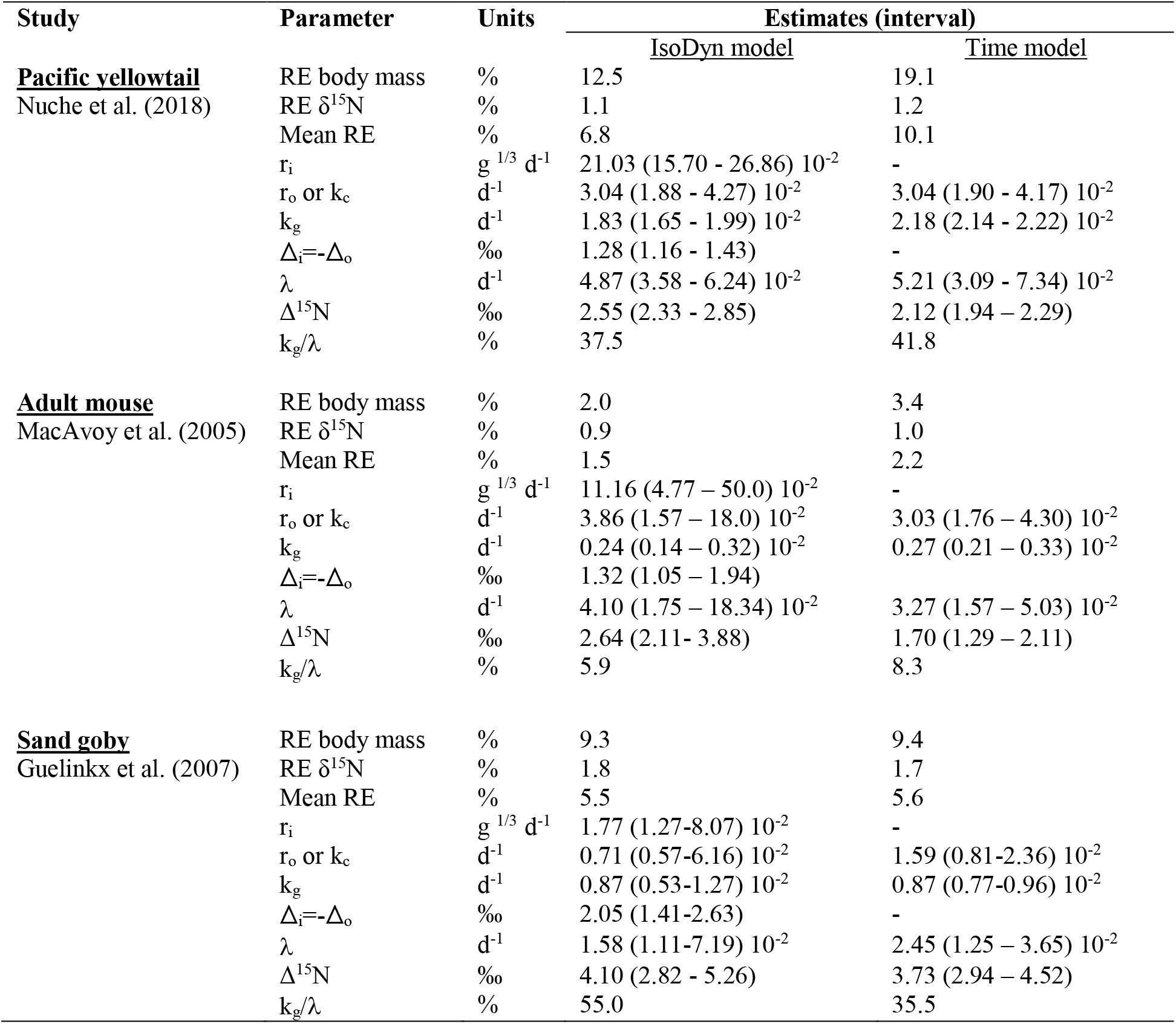
Parameter estimation (best estimates and interval estimates) of the two isotopic incorporation approaches: (1) the time model assuming exponential growth, TIM; (2) the IsoDyn model assuming asymptotic growth patterns with an allometric coefficient β=2/3 for the three case studies. Interval estimates are 95% confidence interval for TIM, and 2.5% and 97.5% quantiles for Isodyn as parameter interval density distribution did not follow a normal distribution. Relative Error (RE, %) between observations and predictions for the body mass and the δ^15^N values, and the mean RE between the two latter. Specific parameters of Isodyn model are r_i_ (rate of gains or assimilation), r_o_ (rate of losses or excretion equivalent to the catabolic rate k_c_), Δ_i_ and Δ_o_ (isotopic discrimination on gains and losses respectively) related to eq(8). In Isodyn model, the isotopic incorporation rate (λ), the specific growth rate (k_g_) and the asymptotic trophic discrimination factor (Δ^15^N) are calculated following eq (7), eq (10), and as Δ_i_ - Δ_o_ respectively. Specific parameters of the time model are λ, Δ^15^N, k_g_ and k_c_ related to eq (1 to 4). Standard propagation of error formulae were used to estimate interval of parameters not directly estimated from the fitting methods

The estimated parameters were of the same order of magnitude for both models but in some cases they differed substantially, especially for λ, k_g_ and k_c_ (Table 2). The estimates of Δ values were roughly the same in both models, although slightly higher in the case of Isodyn model (10 to 20% for sand goby and Pacific yellowtail, respectively). k_g_ were identical in the case of sand goby and smaller for the two other species which body mass dynamics deviated from the exponential pattern. λ estimates were comparable for the first two species (mouse and Pacific yellowtail) but it was 60% smaller using IsoDyn model compared to the time model for the sand goby. As a result, the proportion of k_g_ explaining λ differed noticeably from one case to another. In mouse and Pacific yellowtail, k_g_/λ ratios were 10 to 20% lower respectively in IsoDyn estimates, whereas it was 60% higher for the sand goby. As for the specific estimates of the IsoDyn model, r_i_ ranked according to the maximum body mass of the species (Pacific yellowtail > mouse > sand goby) and r_o_ was higher for mouse (endotherm species). The predicted maximum body masses (as calculated with eq(7)) were 331.5 g, 23.5 g and 15.5 g for Pacific yellowtail, mouse and sand goby respectively. Isotopic discrimination on assimilation or excretion rates (Δ_i_ and Δ_o_ respectively) ranged from 0.8 to 2.1 ‰ The strong interplay of growth and isotopic incorporation dynamics in IsoDyn model offer new perspectives in the interpretation of λ and Δ and a more consistent evaluation of the contribution of growth and catabolic rates in λ.

### Case study: calibration using the sequential parameter estimation

To determine the suitability of IsoDyn model when intra-specific variations of r_i_ and r_o_ occurs due to different ration levels, we re-analyzed the data from Gaye Siesseger et al. (2004). In this DSE, the growth of Common carp (*Cyprinus carpio*) was manipulated by changing food availability through different feeding levels. We estimated parameters in a sequential approach because dynamics of body mass were restricted to start and end values preventing a reliable calibration of r_i_ and r_o_. First, we used parameters from the DEB “Add My Pet” data set to calibrate the model (see supplementary material 2 for more details), and then adjusted the scaled functional response (f), an Holling type II function ranging from 0 to 1, that controls the rates of body mass gains (r_i_) and of body mass losses (r_o_) to fit the observations of body mass change over time (Fig. 5). The f values were estimated to be 0.16, 0.29, 0.53, 0.82 from the lowest feeding ration levels to the highest ones (Table 2). This allowed estimating values of r_i_ and r_o_ for the four treatments according to ESM2 (Table 3). Then, independent values of Δ_i_ and Δ_o_ were simultaneously fitted using δ^15^N_b_-δ^15^N_d_ values and giving Δ_i_ = 1.08‰ and Δ_o_ = −1.32‰. The model not only predicted the qualitative decrease of δ^15^N_b_-δ^15^N_d_ values with k_g_ (Fig. 5) but, in addition, it yielded a very good quantitative fit to the data.

**Fig. 5.**
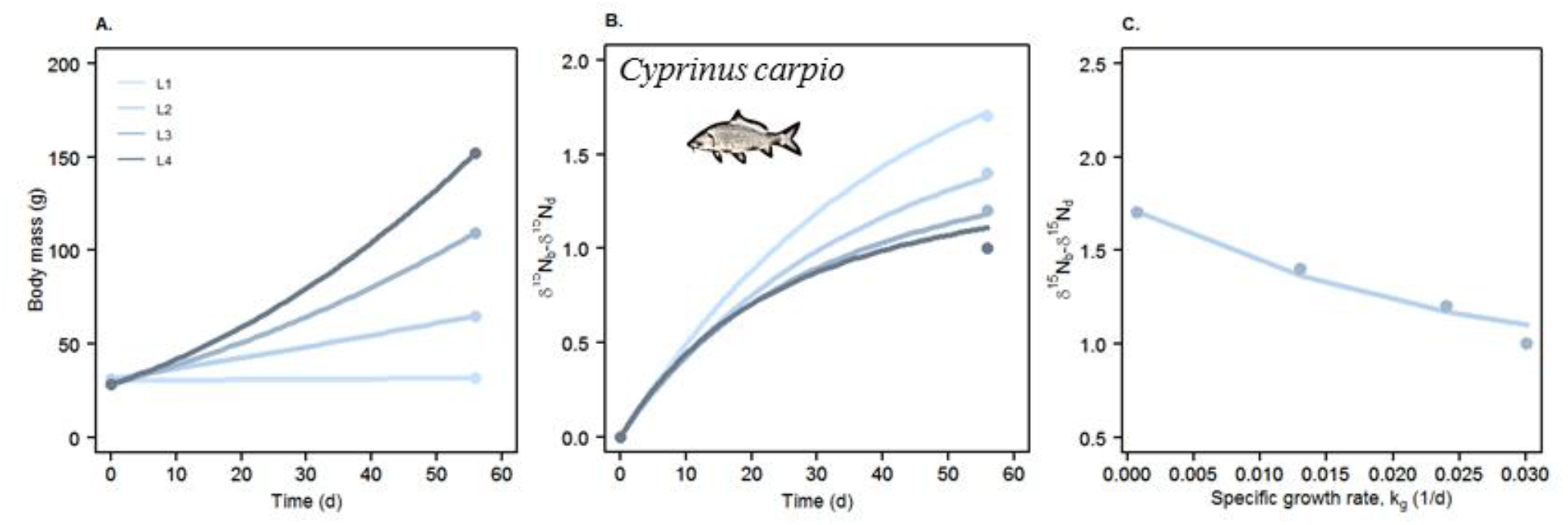
Changes in body mass (A) and δ^15^N_b_ values of whole body minus δ^15^N_d_ values of the diet (B) in common carp (*Cyprinus carpio*) diet-shifted to a new diet and fed four different feeding ration levels (L1, L2, L3, L4; Gaye-Siesseger et al. 2004). Closed circles are observations and lines are predictions. C) represents δ^15^N_b_-δ^15^N_d_ for each diet and hence for each mass specific growth rates

**Table 3.** Estimates of scaled functional response (f), a Holling type II functional response ranging from 0 to 1, of the von Bertalanffy model as predicted by DEB theory (see supplementary material 2 for details) and related r_i_ (rate of gains i.e. assimilation) and r_o_ (rate of losses i.e. excretion) values from the Common carp case study (Gaye-Siesseger et al. 2004). Fish were fed at four feeding ration levels (L) from the lowest feeding ration levels to the highest ones (1 to 4)

| Feeding ration level | $f$ (unitless) | $r_i$ ($\text{g}^{1/3} \text{d}^{-1}$) | $r_o$ ($\text{d}^{-1}$) |
| --- | --- | --- | --- |
| L 1 | 0.16 | $7.39 \cdot 10^{-2}$ | $2.27 \cdot 10^{-2}$ |
| L 2 | 0.29 | $9.62 \cdot 10^{-2}$ | $1.37 \cdot 10^{-2}$ |
| L 3 | 0.53 | $12.34 \cdot 10^{-2}$ | $0.78 \cdot 10^{-2}$ |
| L 4 | 0.82 | $14.47 \cdot 10^{-2}$ | $0.52 \cdot 10^{-2}$ |

## Discussion

### The IsoDyn model links growth and dynamics of isotopes incorporation

Existing dynamic models of isotope incorporation (DIIM) can be ranked in a continuum that spans from simple phenomenological models with few parameters (e.g. Fry and Arnold 1982; Tieszen et al., 1983) to complex mechanistic models with many parameters (e.g. Pecquerie et al. 2010; Poupin et al. 2014). The first models developed were function of either body mass (Fry et al., 1982) or time (Tieszen et al., 1983). Each of these models was then improved later (Carleton and Martinez del Rio, 2010, Heisslein et al., 1993 respectively) in order to partition the λ into two components, k_g_ and k_c_. As these improved models need an independent estimation of k_g_, they do not explicitly connect the underlying mechanisms common to both isotopes and body mass dynamics such as rate of mass gains (r_i_) and rate of mass losses (r_o_). Further, parameters from previous models are constant with time and body mass dynamics are restricted to the exponential or the steady state cases. However, they are simple to use and describe experimental available data sets well in most cases, and have interpretable parameters but they are limited in that they can hide important details of the factors that shape the process of isotope incorporation in a dynamic way.

IsoDyn model renders λ dynamic by considering common and explicit parameters (r_i_, r_o_ and β) to both δ^15^N and body mass dynamics, and offers the possibility to reproduce different growth patterns over the organism life span. This highlights a first important feature of the new model over the previous ones. A second important property of IsoDyn is the possible temporally variable trophic discrimination factor (Δ^15^N) due to its interaction with growth. Our model allows for this interaction thanks to two fluxes of which the flux of gains is allometrically related to body mass, plus that each of the fluxes being associated with a discrimination value. Actually, Δ^15^N varies over time along with growth only if the discrimination linked to body mass losses (Δ_o_) is different from zero (and most probably below zero). Interaction between Δ^15^N and growth was evidenced in experimental results (e.g. Lefebvre and Dubois 2016; Gorokhova 2018) and predicted by earlier mechanistic models but for the exponential case only (Olive et al. 2003; Martinez Del Rio et al. 2005). This interaction is absent in the conventional models developed earlier. IsoDyn model has the time model as a special case (i.e. when λ and Δ^15^N are constants, β=1 and Δ_o_=0) and shares with it ease of computation and analytical tractability.

Our model provides also a new link between models that describe isotopic incorporation phenomenologically and those that incorporate more mechanistic details. Pecquerie et al. (2010) and Emmery et al. (2011) applied dynamic energy budget theory (DEB) to clarify the processes that determine both λ and Δ^15^N values in an approach called Dynamic Isotope Budget modeling (DIB). Unlike our model, DIB models cannot be summarized simply as they are assumption-rich (Pecquerie et al. 2010). DIB recognizes the dynamic dependence of isotope incorporation dynamics on body mass and growth (Emmery et al. 2011). The results of DIB are consistent with our simpler mechanistic model. However, DIB is a two sequential compartments and three fluxes model at least, and then Δ^15^N is not only explained by the isotopic discrimination on fluxes but the proportion of the two sequential compartments that account for the body mass of organisms (i.e. reserve and structure compartments, Pecquerie et al. 2010; Lefebvre and Dubois 2016). In another approach, Poupin et al. (2014) developed a detailed mechanistic multi-compartment model of nitrogen pool and fluxes (21 compartments and 49 fluxes) on adult rat. They showed for instance a deviation from optima in food quality or quantity led to an increase of Δ^15^N at the whole body scale. Unlike IsoDyn model, DIB and the multi-compartment model demand detailed parameterization. The model that we describe shares some of the powerful characteristics of DIB or the multi-compartmental model while making it consistent with the mass-balance models more widely used by isotopic ecologists.

### Parameterizing the model and the experiments that we need

Because growth is a central feature of an animal’s ecological traits, IsoDyn model allows linking patterns of isotopic incorporation and trophic discrimination factor with the biology of animal life histories. This true link between growth and isotope incorporation offers several possibilities regarding the parametrization of our model in simultaneous and sequential estimations: calibration of body mass dynamics and of dynamics of isotope incorporation could insight from each other.

The simultaneous approach allows strengthening the parametrization by coupling body mass dynamics and isotope incorporation dynamics into a single calibration procedure. The limit of this procedure stands in the number of parameters to be estimated since the higher the number of parameters, the bigger the problem of multiple local minima in the minimisation of the cost function. Specifying some parameters is then necessary to relax this problem. This was performed by assuming a known β (here β=2/3), and that the two isotopic discrimination were equal in absolute values (Δ_i_=-Δ_o_). Fittings were then very good. The rate of body mass gains (r_i_) ranked with maximal body size and this is coherent with metabolic theories (West et al. 2001; Kooijman 2010). Estimates of r_i_ and r_o_ allow to predict the maximum body masses that can be reached by the three species using eq(6). The maximal body mass was correctly estimated for mouse (23.5 g vs 25 g) and sand goby 15.5 g vs 7 g) but was underestimated for Pacific yellow tail (331.5 g vs 193 kg) probably due to sub-optimal experimental conditions for this large and migratory fish species.

One way to improve the estimation of IsoDyn model parameters in the simultaneous estimation is to perform DSE with different conditions of growth for the same species fed with the same diet with measurement of body mass dynamics in parallel. It can be done performing either DSEs at different life stages of the same species with the same diet to satiation, or DSEs at different food rations at one life stage when growth is still significant. To our best knowledge, the first case has not been reported yet in literature. The second one is rare and the body mass dynamics with an adequate time resolution were not reported (e.g. Gaye Siesseger et al. 2004; Lefebvre and Dubois 2016; Gorokhova 2018;). Unfortunately, most of DSEs reporting both body mass and δ^15^N dynamics used diets that differ in quality and are fed to satiation (Nahon et al. 2020), and this leads to different growth rates but possibly confounding results with additional sources of λ and Δ^15^N variations (e.g. diet type, mode of nitrogen excretion, etc…).

In the sequential parameter estimation, the model’s simplicity allows ready parameterization with available estimates of r_i_, r_o_ and β (Common carp case study). Because our new model incorporates widely used individual growth models, it links isotopic ecology with large bodies of data (Ontogenetic growth model, West et al. 2001; Hou et al. 2011, von Bertalanffy models, including DEB’s “Add My Pet” data base and other growth rate data available for a large number of animals, Marques et al. 2018). The two approaches can explore the effect of food restrictions on body mass dynamics (Kearney 2020). Once the individual growth model is parameterized, calibrations of isotopic discriminations on flux of body mass gains and losses (Δ_i_ and Δ_o_ respectively) can be easily performed. Results from the Common carp case study have emphasized that Δ_i_ was a bit lower than Δ_o_. The latter is probably the main driver of Δ^15^N enrichment in animal tissues (Poupin et al. 2014).

The range of values of Δ_i_ and Δ_o_ can be predicted from the relationship between Δ^15^N and k_g_: When k_g_ is high for a given species Δ^15^N is mostly explained by Δ_i_ whereas when k_g_ approaches 0, Δ^15^N equals Δ_i_-Δ_o_. For example, Δ^15^N varied between 2 and 4‰ depending on k_g_ in mysids (Gorokhova 2018), from 3 to 9‰ in invertebrates (Lefebvre and Dubois 2016), from 2 to 5‰ in a bivalve (Emmery et al., 2011) from 1 to 1.7‰ in the Common carp case study (Gaye-Siesseger et al. 2004). From these ranges, one can predict that the Δ_o_ values are probably higher that Δ_i_ values in general. Generalizing the calibration of IsoDyn model on DSEs would help determine the range of the Δ_o_ and Δ_i_ values using meta-analyses on some particular taxons. Finally, a common problem in the interpretation of isotopic data from studies is that the family of eq(1) needs a DSE data set with a clear shift and a clear asymptote to relax as much as possible the co-variation of λ and the asymptotic value (δ_∞_ needed to estimate Δ^15^N) and their uncertainties. Our model relaxes the necessity of perfect DSEs since the calibration can be sequential. DSEs (e.g. Logan and Lutcavage 2010) that provide limited information for λ and Δ^15^N could be then exploited with the IsoDyn model used as an alternative.

### Implications of the IsoDyn model for isotopic ecology

With all its simplifying assumptions, the IsoDyn model represents significant progress. In particular, it offers new perspectives in understanding the variabilities of λ and Δ^15^N values, two critical variables for the interpretation of isotopic data (Martinez del Rio et al. 2012). Vander Zanden et al (2015) or Thomas et al. (2015) constructed allometric relationships that relate λ values with body size and several authors have summarized data on Δ^15^N and searched for the potential causes for its variation (Vanderklift and Ponsard 2003; Caut et al. 2009; Healy et al. 2018). The allometric studies of Vander Zanden et al. (2015) and Thomas et al. (2015) verified the prediction that λ varies as an allometric function of body mass (Martinez del Rio et al., 2009). Although these relationships are in broad agreement with predictions, they have large residual variation that limits precise estimation. We hypothesize that some of this variation can be explained by growth, the factor identified by IsoDyn model as a major determinant of λ and Δ^15^N.

By necessity, these large comparative data sets gloss over the characteristics of the animals that might generate variation in λ and Δ^15^N due to growth. For example, the vast majority of the estimates of λ and Δ^15^N on endotherms with determinate growth like birds and mammals are done on fully-grown adults. The same is the case of measurements of small invertebrates that reach asymptotic body mass in a short time. In contrast, experiments on ectotherms with indeterminate growth such as fish, amphibians, and reptiles are done in growing juvenile animals. This growth effect may explain why Δ^15^N mean values in ectotherms are slightly lower than the ones on endotherms (Caut et al. 2009). Re-analysing results of these meta-analyses using the IsoDyn model would be an interesting perspective. Further, we identified areas in which its application can solve long-standing questions to merge isotopic ecology and trophic ecology more seamlessly: the reconstruction of diet, the interpretation of “isotopic niches” and the determination of trophic level and food web structure.

Stable isotopes are very often used within mixing models to estimate the proportions of dietary items with contrasting isotopic values into animal diets at species (Layman et al. 2012) and food web level (see Kadoya et al. 2012). Indeed the use of mixing models to estimate diet proportions has increased exponentially over the last years (as referred to in the citation dynamics of Parnell et al. 2010 paper). The mixing models used for this purpose require estimates of Δ^15^N and assumed isotopic equilibrium between diet and consumers. Relaxing the isotopic equilibrium assumption has been the concern of several studies with different prospects but in which the Isodyn model may help to quantify the parameter values. Phillips et al. (2014) recommended to carefully consider the time period over which the putative food sources have to be sampled to back calculate diet using mixing-models. Actually, this time period relies on λ(Thomas and Crowther 2015). Stock and Semmens (2016) integrated a new component in their mixing models by accounting for the variation in consumption rate between individuals of a population. The rate of body mass gains (r_i_) is a proxy of this consumption rate.

Hertz et al. (2016) evidenced that λ is a critical parameter when modelling ontogenetic diet shifts. They typically modified a growth incorporation model in which λ vary with the body mass increase (Fry and Arnold 1982), but they kept constant the contribution of k_g_ and k_c_ while it is variable in the IsoDyn model. Finally, many ecologists analyse muscle for large species or whole body for small species, those working on endotherms use blood, and paleontologists are constrained to the analysis of bone and collagen. These tissues have widely different λ values (Thomas and Crowther 2015) and can have different Δ^15^N within an organism (Vanderklift and Ponsard 2003). A multi-compartment extension of the Isodyn model might allow predicting the magnitude of Δ^15^N values among tissues and the effect of growth on these values. Building a multi-compartment extension of the IsoDyn model has both computational and empirical challenges. Martínez del Rio and Andreson-Sprecher (2008) described how to arrange several compartments in parallel or sequentially (or a mix of both as in DIB for adults) but the model used assumes steady state or exponential growth and like all conventional models it assumes no dynamic pattern for λ and Δ^15^N. Unlike Poupin et al.’s (2014) model which assumes that the animals are not growing and hence allows using a system of linear differential equations, the multi-compartment Isodyn model is non-linear and hence is computationally more complex. Furthermore, the model requires empirical data of changes in fluxes among compartments that can vary in relative size during development or not. Challenging as they will be, these models are needed to estimate observed differences in both λ and Δ^15^N in different organs.

Our model suggests that differences in k_g_ can distort the geometry of isotopic niches beyond the frequency of diet change (Yeakel et al. 2016). The characteristics of the space occupied by individuals, populations, and by species assemblages in isotopic space are often used to interpret trophic structure (Shipley and Matich 2020). For example, the area of standard ellipses (and other metrics of extent of occupancy of isotopic space, Layman et al. 2012) is often used to assess variation in resource use (Parnell et al. 2013). Gorokhova (2018) demonstrated experimentally that the characteristics (as assessed by commonly used metrics) of the “isotopic niches” were dependent on growth (and hence on feeding regime) in Mysid shrimp (*Neomysis integer*) fed on the same food but different rations. In accordance with the results of IsoDyn model, she found lower Δ^15^N in animals fed at high rations and hence growing more rapidly. Through its effects on λ and Δ^15^N, k_g_ can change the position and variance (as measured by area occupied in isotopic space) of isotopic niches all the more so as growth is time dependent. The interpretation of isotopic patterns must be informed by the mechanisms that shape them, including growth rate.

The value of Δ^15^N is not only used in mixing models applied to determine diet composition. It is also often used to estimate an animal’s trophic position in a food web (Post 2002; Quezada-Romegialli et al. 2018). An extension of this application is the use of the interspecific range of Δ^15^N values in assemblages of consumers to estimate the length of a food chain (Vander Zanden and Fetzer 2007). The prediction of our model adds a note of caution to the interpretation of the use of stable isotopes as estimates of trophic position and food-chain length, but opens the opportunity to make these measurements more accurate. Villamarin et al. (2018) identified a clear mismatch between trophic position estimated from Δ^15^N measurements and diet in crocodiles. This mismatch was largely explained by a decrease in Δ^15^N with k_g_ consistent with the predictions of the IsoDyn model (Villamarín et al. 2018). The often reported positive correlation between Δ^15^N and body size in fishes (e.g. Nakazawa et al. 2010) that is attributed to upwards shifts in trophic position might have to be reconsidered in light of declining growth rates (and hence Δ^15^N values) with size predicted by our model. This could also have additional unsuspected consequences when scaling δ^15^N values and trophic level (Hussey et al. 2014).

So far, patterns of occupancy in isotopic space are used to infer the ecological characteristics of individuals, populations, and food wed structure. Our model suggests that patterns in measured isotopic values are not only the result of a one-way translation of resource use into isotopic value. They are the dynamic outcome of not only how animals use resources, but of the tempo and fidelity of isotopic incorporation. These are shaped by the mechanisms by which animals incorporate and dispose materials into their tissues. We believe that incorporating these mechanisms into dynamic models can transform isotopic ecology from a descriptive into a more dynamic process-based discipline. Recent studies advocated for the use of simulation modelling to predict stable isotope ratios using mechanistic processes (e.g. Flynn et al. 2018; Trueman et al. 2019). Isodyn model can be an element of these models, and hence can be a further step in the direction of a mechanistic process-based isotopic ecology.

## Supporting information

Electronic Supplementary Material 1

Electronic Supplementary Material 2

## Declaration

### Ethical approval

This article does not contain any studies with human participants or animals performed by any of the authors

