## Supplementary material for "Individual growth models support the quantification of isotope incorporation rate, trophic discrimination and their interactions": Electronic Supplementary Material 1

Development of the IsoDyn model

Our model assumes that an individual grows following the rule:

$$\frac{dW}{dt} = r_i W^{\beta} - r_o W \qquad \text{eq(S1.1)}$$

where W is the body mass of the individual and $r_i$ and $r_o$ are constants that govern the rates of mass gains and losses respectively and β is the allometric coefficient. The value of the exponent β depends on whether the flux of mass gain, $r_i W^{\beta}$ is dependent on surface area as proposed by von Bertalanffy (1957), in which case β=2/3, or to the exponent proposed by the ontogenetic growth model (West et. al. 2001), in which case β=3/4. For our purposes, the value of β can be simply assumed to range between 0.5 and 1 (Dodds et al., 2001) and, when needed, estimated empirically. A general solution for asymptotic growth models (i.e. with asymptotic body mass or length) is developed below.

Define $U_t = W_t^{1-\beta}$, with

$$\frac{dU}{dt} = (1-\beta)W^{-\beta}\frac{dW}{dt} = (1-\beta)W^{-\beta}\left(r_i W^{\beta} - r_o W\right) \qquad \text{eq(S1.2)}$$

and

$$\frac{dU}{dt} = (1-\beta)r_i - (1-\beta)r_o U \qquad \text{eq(S1.3)}$$

This equation is linear with solution:

$$U_t = U_0 \mathrm{e}^{-r_o(1-\beta)t} + \frac{r_i}{r_o}\left(1 - \mathrm{e}^{-r_o(1-\beta)t}\right) \qquad \text{eq(S1.4)}$$

but because $U_t = W_t^{1-\beta}$

$$W_t^{1-\beta} = W_0^{1-\beta}\mathrm{e}^{-r_o(1-\beta)t} + \frac{r_i}{r_o}\left(1 - \mathrm{e}^{-r_o(1-\beta)t}\right) \qquad \text{eq(S1.5)}$$

and

$$W_t = \left\{W_\infty^{1-\beta} + \left(W_0^{1-\beta} - W_\infty^{1-\beta}\right)\mathrm{e}^{-r_o(1-\beta)t}\right\}^{\frac{1}{1-\beta}} \qquad \text{eq(S1.6)}$$

with

$$\lim_{t\to\infty} W_t = W_\infty = \left(\frac{r_i}{r_o}\right)^{\frac{1}{1-\beta}} \qquad \text{eq(S1.7)}$$

And with

$$k_g = \frac{1}{W}\frac{dW}{dt} = r_i W_t^{1-\beta} - r_o \qquad \text{eq(S1.8)}$$

Assuming that N pool is a fraction $p_N$ of W with N= $p_N$ W, this gives from eq(S1):

$$\frac{dN}{dt} = r_i p_N^{1-\beta} N^{\beta} - r_o N \qquad \text{eq (S1.9)}$$

Then

$r_{iN} = r_i p_N^{1-\beta}$ and $r_{oN} = r_o$ with $r_{iN}$ and $r_{oN}$ the constants that govern the rates of N gains and N losses respectively.

The pool of nitrogen (N) is the sum of nitrogen light isotope ($N_l$) and nitrogen heavy isotope ($N_h$) and $N_h$ follows:

$$\frac{dN_h}{dt} = (f_d + \epsilon_i) r_{iN} N^{\beta} - (f_b + \epsilon_o) r_{oN} N \qquad \text{eq (S1.10)}$$

where $f_d$ and $f_b$ are the heavy isotope fractions of the diet and body of the consumer, respectively. The constants $\in_i$ and $\in_o$ represent isotopic discrimination during nitrogen gains (i.e. assimilation) and nitrogen losses (i.e. excretion or respiration in the case of carbon), respectively $\in_i = f_{ass} - f_d$ and $\in_o = f_{excr} - f_b$, where $f_{ass}$ and $f_{excr}$ are the fraction of heavy isotopes assimilated and excreted, respectively, and $\in_i > 0$ and $\in_o < 0$). By definition, $f_b = \frac{N_h}{N}$, therefore Eq(S9) and Eq(S10) can be combined to yield the change in the fraction of the heavy isotope in the animal's body as:

$$\frac{df_b}{dt} = \frac{1}{N}\left(\frac{dN_h}{dt} - \frac{dN}{dt} f_b\right) = r_{iN} N^{\beta-1}(f_d - f_b + \epsilon_i) - \epsilon_o r_{oN} \qquad \text{eq(S1.11)}$$

Throughout, we model changes in isotopic composition through time using fractions of the heavy element, but we present our results in the traditional $\delta$ notation using the approximation:

$$f_i \approx R_{std}\left(\frac{\delta^{15}N}{1000} + 1\right) \qquad \text{eq(S1.12)}$$

where $f_i$ represents the fraction of the heavy element in each of the components modelled ($f_d$ and $f_b$ for diet and body respectively). For sake of simplicity, we remove the "b" subscript in the following equations. In the same vein, the constants $\in_i$ and $\in_o$ are noted $\Delta_i$ and $\Delta_o$ and the trophic discrimination factor $\Delta$ becomes $\Delta^{15}N$. Further, as $r_{iN} = r_i p_N^{1-\beta}$ and $N = p_N W$, then eq(S11) becomes:

$$\frac{d\delta^{15}N}{dt} = r_i W^{\beta-1}(\delta^{15}N_d - \delta^{15}N + \Delta_i) - \Delta_o r_o \qquad \text{eq(S1.13)}$$

Eq(S13) is a linear differential equation without a general analytical solution when β is lower than 1 (i.e. for asymptotic growth models such as the von Bertalanffy one). However, for a given and constant body mass, Eq(S13) has the solution :

$$\delta^{15}N_t = e^{-r_i W_t^{\beta-1} t}\left\{cst - \left(\delta^{15}N_d - \Delta_i + \frac{\Delta_o r_o}{r_i W_t^{\beta-1}}\right) e^{-r_i W_t^{\beta-1} t}\right\} \qquad \text{eq(S1.14)}$$

At time t=0

$$\delta^{15}N_0 = cst - \left(\delta^{15}N_d - \Delta_i + \frac{\Delta_o r_o}{r_i W_t^{\beta-1}}\right) \qquad \text{eq(S1.15)}$$

Giving

$$cst = \delta^{15}N_0 + \left(\delta^{15}N_d - \Delta_i + \frac{\Delta_o r_o}{r_i W_t^{\beta-1}}\right) \qquad \text{eq(S1.16)}$$

Assuming that when t→∞, we have :

$$\delta^{15}N_\infty = \left(\delta^{15}N_d - \Delta_i + \frac{\Delta_o r_o}{r_i W_t^{\beta-1}}\right) \qquad \text{eq(S1.17)}$$

Then eq(S14) becomes

$$\delta^{15}N_t = \delta^{15}N_\infty - (\delta^{15}N_\infty - \delta^{15}N_0)e^{-r_i W_t^{\beta-1} t} \qquad \text{eq(S1.18)}$$

Eq(S13) can be solved numerically using numerical integration algorithms. Alternatively, we can obtain an accurate approximation using Eq(S18) in an iteration procedure for small time interval (small dt) using a discrete approximation

$$\delta^{15}N_{t+1} = \delta^{15}N_{\infty} - (\delta^{15}N_{\infty} - \delta^{15}N_t)e^{-r_i W_t^{\beta-1} dt} \qquad \text{eq(S1.19)}$$

Special case 1: steady state growth

If the organism is at steady state at $W_\infty$ (i.e. the case of adult endotherms), $r_i W_\infty^{\beta} - r_o W_\infty = 0$, and $\frac{r_i}{r_o} = W_\infty^{1-\beta}$ giving $r_i = W_\infty^{1-\beta} r_o$ . This further gives $\lambda = r_o$ and $\Delta^{15}N = \Delta_i - \Delta_o$.

In this case the solution to eq(S13) becomes:

$$\delta^{15}N = \delta^{15}N_d + \Delta_i - \Delta_o - (\delta^{15}N_d + \Delta_i - \Delta_o - \delta^{15}N_0)e^{-r_o t} \qquad \text{eq(S1.20)}$$

Our model predicts that in non-growing organisms $\Delta^{15}N$ will not be dependent on λ . Consequently, $\Delta^{15}N$ is constant and at its maximal value.

Special case 2: exponential growth

If the organism is growing exponentially, β=1 and Eq(S1) reduces to:

$$\frac{dW}{dt} = (r_i - r_o)W \qquad \text{eq(S1.21)}$$

and the solution to eq(S1) is still the eq(S18) but with $\delta^{15}N_\infty = \delta^{15}N_d + \Delta_i - \frac{\Delta_o r_o}{r_i}$, and $\lambda = r_i$. Then, $\Delta^{15}N = \Delta_i - \frac{\Delta_o r_o}{r_i}$ and is a decreasing function of λ (i.e. $r_i$) and an increasing function of $\Delta_o$. Also, because λ is a linearly increasing function of mass-specific growth ($\lambda = k_g + r_o$), the model predicts that $\lambda$ increases with mass-specific growth rate. Thus, the relationship between $\Delta^{15}N$ and $\lambda$ is again mediated by growth if $\Delta_o$ differs from 0‰. Note that if $\Delta_o$ =0‰ and $\Delta_i$ >0‰ eq(S18) becomes the time model (eq(1)). The exponential phase is in many cases an early phase of asymptotic growth models (Kearney 2020). The true exponential case with β=1 will not be considered further.
