## Supplementary material for "Individual growth models support the quantification of isotope incorporation rate, trophic discrimination and their interactions": Electronic Supplementary Material 2

Link between Von Bertalanffy growth model, DEB and IsoDyn

The dynamic energy and mass budget (DEB) theory ambitions to model the quantitative aspects of the organization of metabolism of any living organisms by depicting common rules of allocations and processes at the organism level (Sousa et al., 2008). DEB model consists in a system of three ordinary differential equations following first order dynamics. Assimilated products first enter a reserve pool (E) which is then mobilized to fuel two pathways following the κ rule: a fixed κ fraction is allocated to perform growth of the structural volume (V) and its maintenance and the remaining fraction (1-κ) is available for maturity (i.e. increase of complexity and its maintenance) and reproduction (Van der Meer, 2006). The DEB theory describes the energy flows within an organism for a given food level and at a reference temperature $T_{ref}$. Rates are temperature corrected $T_{cor}$ using the following equation:

$$\dot{k}(T) = \dot{k}_1 T_{cor} = \dot{k}_1 e^{\left(\frac{T_A}{T_{ref}} - \frac{T_A}{T}\right)} \qquad \text{eq(S2.1)}$$

where $T_A$ is the Arrhenius temperature (in °K), $\dot{k}_1$ the rate of interest at the reference temperature, $T_{ref}$ , and $\dot{k}(T)$ the rate of interest at temperature T (in °K).

The assimilation rate is described by an hyperbolic functional response called the scaled functional response, f, a Michaelis-Menten function also named the Holling type II functional response. f is a limiting function that varies between 0 (i.e. starvation) and 1 (i.e. satiety) depending on the substrate concentration (i.e. food availability).

The standard DEB model considers an isomorphic organism assuming that food assimilation rate is dependent on surface area (i.e. body volume raised to the power 2/3) and maintenance is proportional to body volume. For a given reserve density (i.e. for a given food density, f), DEB model explains the so-called von Bertalanffy growth model.

$$\frac{dL}{dt} = \dot{r}_B (L_\infty - L) \qquad \text{eq(S2.2)}$$

with L the shape corrected structural length, $L_\infty$ the asymptotic structural length and $\dot{r}_B$ the von Bertalanffy growth rate.

In DEB theory, body mass (W) is the sum of weight of the reserve compartment plus the one of the structural volume (V or $L^3$ assuming the density of wet structure is equal to 1 $g.cm^{-3}$). Wet weight of the reserve compartment depends on the structural length, the scaled reserve density (equal to f at constant food availability), and the contribution of reserve to body mass (cbm), giving:

$$W = L^3 + L^3 f\ cbm = L^3(1 + f\ cbm) \qquad \text{eq(S2.3)}$$

Assuming that $W = aL^3$ and $L = \frac{W^{\frac{1}{3}}}{a^{\frac{1}{3}}}$ or $L^2 = \frac{W^{\frac{2}{3}}}{a^{\frac{2}{3}}}$ with a equal to (1+f cbm)

$$\frac{dW}{dt} = \frac{dW}{dL}\frac{dL}{dt} = 3aL^2 \dot{r}_B (L_\infty - L) \qquad \text{eq(S2.4)}$$

$$\frac{dW}{dt} = 3a\frac{W^{\frac{2}{3}}}{a^{\frac{2}{3}}}\dot{r}_B\left(\frac{W_\infty^{\frac{1}{3}}}{a^{\frac{1}{3}}} - \frac{W^{\frac{1}{3}}}{a^{\frac{1}{3}}}\right) \qquad \text{eq(S2.5)}$$

$$\frac{dW}{dt} = 3\dot{r}_B W_\infty^{\frac{1}{3}} W^{\frac{2}{3}} - 3\dot{r}_B W \qquad \text{eq(S2.6)}$$

with $W_\infty$ the asymptotic body mass.

In DEB theory, the asymptotic structural length is a function of food availability (f), the ultimate structural length $L_{max}$ and a correction factor $s_M$ which accounts for a metabolic acceleration at early ontogenetic phase in some species. In standard DEB model $s_M$ =1 and in "abj" typified model $s_M$ is the ratio between length at juvenile stage and length at birth stage. Then $W_\infty$ is equal to:

$$W_\infty = (f L_{max} s_M)^3(1 + f\ cbm) \qquad \text{eq(S2.7)}$$

In DEB theory, the von Bertalanffy growth rate ($\dot{r}_B$) follows:

$$\dot{r}_B = \frac{1}{3}\frac{k_M}{1+\frac{f}{g}} T_{cor} \qquad \text{eq(S2.8)}$$

with $k_M$ the somatic maintenance rate coefficient ($d^{-1}$), f the scaled functional response (unitless), g the energy investment ratio (i.e. the costs of an increase in body mass relative to the maximum potentially available energy for growth plus maintenance; unitless) and $T_{cor}$ the temperature correction.

Following our notation, the Von Bertalanffy growth model for body mass is written as:

$$\frac{dW}{dt} = r_i W^{\frac{2}{3}} - r_o W \qquad \text{eq(S2.9)}$$

Then

$$r_o = 3\dot{r}_B = \frac{k_M}{1+\frac{f}{g}} T_{cor} \qquad \text{eq(S2.10)}$$

And

$$r_i = 3\dot{r}_B W_\infty^{\frac{1}{3}} = \frac{k_M}{1+\frac{f}{g}} T_{cor} (f L_{max} s_M)^3 (1 + f\ cbm) \qquad \text{eq(S2.11)}$$

The requested DEB parameters and their values can be retrieved from the Add my pet collection (Marques et al., 2018) for more than 2000 species to date. An example is given in the below table S1 for Common carp (*Cyprinus carpio*). For a temperature of 27 °C (300.15 °K), the temperature correction is worth 1.89. Then for a scaled functional response f=1 (i.e. satiety), $r_o$= 0.0043 $d^{-1}$, $W_\infty$ = 47623.34 g and $r_i$ = 0.156 $d^{-1}$.

Table S1. Requested DEB parameters to parameterize the von Bertalanffy model (body mass) for Common carp (*Cyprinus carpio*)

| Parameter name | Symbol | Value | Unit |
|---|---|---|---|
| The shape corrected ultimate structural length | $L_{max}$ | 6.3743 | cm |
| The acceleration factor | $s_M$ | 1.2236 | unitless |
| The somatic maintenance rate coefficient | $k_M$ | 0.0698 | $d^{-1}$ |
| The energy investment ratio | g | 0.0337 | unitless |
| The contribution of reserve to body mass | cbm | 99.37 | unitless |
| The Arrhenius temperature | $T_A$ | 8000 | °K |
| The reference temperature | $T_{ref}$ | 293.15 | °K |

Marques, G.M., Augustine, S., Lika, K., Pecquerie, L., Domingos, T., Kooijman, S.A.L.M., 2018. The AmP project: comparing species on the basis of dynamic energy budget parameters. PLoS Comput. Biol. 14, 1–23. https://doi.org/10.1371/journal.pcbi.1006100

Sousa, T., Domingos, T., Kooijman, S. A. L. M., 2008. From empirical patterns to theory: A formal metabolic theory of life. Phil. Trans. R. Soc. B, 363, 2453-2464. https://doi.org/10.1098/rstb.2007.2230

Van Der Meer, J., 2006. An introduction to Dynamic Energy Budget (DEB) models with special emphasis on parameter estimation. J. Sea Res. 56, 85–102. https://doi.org/10.1016/j.seares.2006.03.001
